# Faster model updating in autism during early sensory processing

**DOI:** 10.1101/2020.09.04.279471

**Authors:** Judith Goris, Senne Braem, Shauni Van Herck, Eliane Deschrijver, Jan R. Wiersema, Bryan Paton, Marcel Brass, Juanita Todd

## Abstract

**Background:** Recent theories of autism propose that a core deficit in autism would be a less context-sensitive weighting of prediction errors. There is also first support for this hypothesis on an early sensory level. However, an open question is whether this decreased context-sensitivity is caused by faster updating of one’s model of the world (i.e. higher weighting of new information), proposed by predictive coding theories, or slower model updating. Here, we differentiated between these two hypotheses by investigating how first impressions shape the mismatch negativity (MMN), reflecting early sensory prediction error processing.

**Methods:** An autism and matched control group (both *n*=27) were compared on the multi-timescale MMN paradigm, in which tones were presented that were either standard (frequently occurring) or deviant (rare), and these roles reversed every block. A well-replicated observation is that the initial model (i.e. the standard and deviant sound in the first block) influences MMN amplitudes in later blocks. If autism is characterized by faster model updating, we hypothesized that their MMN amplitudes would be less influenced by the initial context.

**Results:** We found that MMN responses in the autism group did not differ between the initial deviant and initial standard sounds as they did in the control group.

**Conclusions:** These results show that individuals with autism are less influenced by initial contexts, confirming that autism is characterized by faster updating of sensory models, as proposed by predictive coding accounts of autism.

## Introduction

Autism spectrum disorder (henceforth “autism”) is characterized by persistent deficits in social interaction and communication, and nonsocial symptoms like restricted, repetitive patterns of behavior (American Psychiatric Association, 2013). Ever since autism was first described, researchers have tried to identify one core deficit that can account for this heterogeneous set of characteristics. Recent theories, grounded in the predictive coding framework (Lawson et al., 2014; Van de Cruys et al., 2014; for a review see Palmer et al., 2017), proposed that a core deficit in autism might be a problem in the flexible weighting of prediction errors in autism. In other words, individuals with autism might have difficulties with estimating in which context a surprising event is important and in which it is not, which could explain a broad range of autism characteristics.

According to the predictive coding framework, the brain constantly makes predictions or “models” about the world and processes all incoming sensory information in comparison to those predictions (Friston, 2010; Rao & Ballard, 1999). The difference between incoming information and predictions is encoded as a surprise signal called a *prediction error*. The brain then uses this signal to adapt its model of the world. However, not all prediction errors are equally important. Therefore, the brain needs to distinguish between important prediction errors that should be used to update its model, and less important prediction errors that can be ignored. It is this ability to flexibly adjust the weight or importance a person gives to their prediction errors that is proposed to be impaired in autism (Lawson et al., 2014; Palmer et al., 2017; Van de Cruys et al., 2014). This single deficit could potentially account for the broad range of autism characteristics, including both social and nonsocial characteristics (for an overview, see Van de Cruys et al., 2014).

Specifically, predictive coding theories of autism propose that people with autism are generally too fast in updating their model of the world based on incoming information, and are therefore less flexible in building broader context-specific models. That is, these theories explain the inflexible weight of prediction errors by a failure to attenuate sensory prediction errors (Lawson et al., 2014) or a general higher weight on prediction errors (Van de Cruys et al., 2014, 2017), leading to new information being incorporated *faster* into the model predictions of people with autism, compared to control participants. However, an inflexible weighting of prediction errors could also be explained by people with autism being *slower* in updating their models of the world. This would mean that older information that is likely irrelevant in the current context is still taken into account. While this is contrary to predictive coding accounts of autism, Lieder and colleagues (2019) showed that participants with autism weighted recent stimuli less heavily in a perceptual decision-making task, and were more influenced by longer-term regularities. Also intuitively, based on autism symptomatology, it can be reasonable to expect that individuals with autism hold on more to information learned in the past, as rigid, inflexible thinking is a key characteristic of autism. Here, we investigate which of these two explanations can account for the previous finding of decreased context-specific prediction errors, by investigating how first impressions influence early sensory processing in autism.

For this purpose, we rely on the multi-timescale paradigm by Todd and colleagues (2011, 2013). In this paradigm, participants are presented with two tones that are either standard (highly probable) or deviant (rare), but these probabilities reverse in alternating blocks of sound that make up the sequences. When exposed to these sequences, sound evoked brain potentials measured from the scalp can be used to assess how the brain rapidly and automatically forms a model of the predicted regularity and produces a prediction error signal, called mismatch negativity (MMN) when a rare deviant tone occurs. By manipulating the length of the blocks, sequences containing slower changing and faster changing alternations can be used to assess how MMN is weighted by different periods of stability. The multi-timescale paradigm has revealed a *primacy bias* such that the initial model (i.e., the respective tones that were the standard and the deviant, in the first block) influences perception in later blocks (Todd et al., 2011, 2013). Specifically, later blocks with reversed contingencies typically show smaller MMN amplitudes (as opposed to those with similar contingencies), especially if they are longer and provide more time to (re)learn one’s model (Todd et al., 2011, 2014). Therefore, using this paradigm, we can investigate the speed with which people update their models. If individuals with autism are faster in updating models of sensory information (and abandoning or unlearning old ones), as suggested by predictive coding theories, we should see a smaller primacy bias (i.e. a reduced influence of initial learning leading to equivalent modulation of MMN amplitudes by the initial model), as we expect them to be less influenced by their older model of the world. However, if autism is characterized by slower model updating, we expect the opposite, namely a larger primacy bias in MMN amplitudes.

## Methods and Materials

### Participants

In total, 30 adults with autism and 30 typically developed (TD) adults participated in the study. Three participants in each group were excluded due to excessive alpha waves in the EEG signal (see below). The final sample size thus consisted of 27 participants with autism (8 female, 5 left-handed, 2 ambidextrous) and 27 TD participants (8 female, 6 left-handed). All participants had a full-scale IQ above 80, and reported to be free of hearing problems. Participants in the TD group were screened to have no reported history of neurological or psychiatric disorders, and have scores below the cutoff on both the Autism Spectrum Quotient (lower than 32) (AQ; Baron-Cohen et al., 2001) and the Social Responsiveness Scale–Adult version (T-score lower than 61) (SRS-A; Constantino, 2002). These are the cutoffs as described in the original questionnaires, meant to screen for autistic traits in an adult population. Age ranged from 23 to 50 years old (*M*=35.63, *SD*=7.54) and did not differ significantly between the two groups, *t*(51.41)=0.83, *p*=.41. All participants gave written informed consent before participation and were financially compensated. The study was approved by the local Ghent University ethics committee.

All adults with autism had received a clinical diagnosis of autism spectrum disorder (*n*=20), Asperger’s syndrome (*n*=6) or autistic disorder (*n*=1), prior to the experiment, by an independent clinician or multidisciplinary team. The diagnoses were verified with the Autism Diagnostic Observational Schedule 2 (ADOS-2) (Lord et al., 2000) Module 4 by a trained researcher using the revised scoring algorithm (Hus & Lord, 2014). In line with earlier autism studies (Deschrijver et al., 2016, 2017; Goris et al., 2018; Magnée et al., 2008), participants with autism were screened to have an ADOS-2 total score of 1 point below the cutoff or higher. Importantly, the results did not change in a statistically significant way when we used the ADOS-2 total score cutoff as the exclusion criterion, which resulted in the exclusion of only one participant.

Intelligence was assessed by using the Kaufman 2 short form Wechsler Adult Intelligence Scale–third edition (WAIS-III). For 12 participants with autism, we used a WAIS-IV or Kaufman 2 short form WAIS-III that was available and completed within the past five years. There was no significant difference in IQ between the two groups, *t*(50.66)=1.63, *p*=.11 (TD group: *M*=118.00, *SD*=12.26; autism group: *M*=112.07, *SD*=14.44).

### Task and Stimuli

The task was an adapted version of the multi-timescale MMN paradigm by Todd and colleagues (2013), which is an auditory oddball paradigm where the regularities defining standard and oddball sounds change over time, in short versus long time windows, see Figure 1.

**Figure 1.**
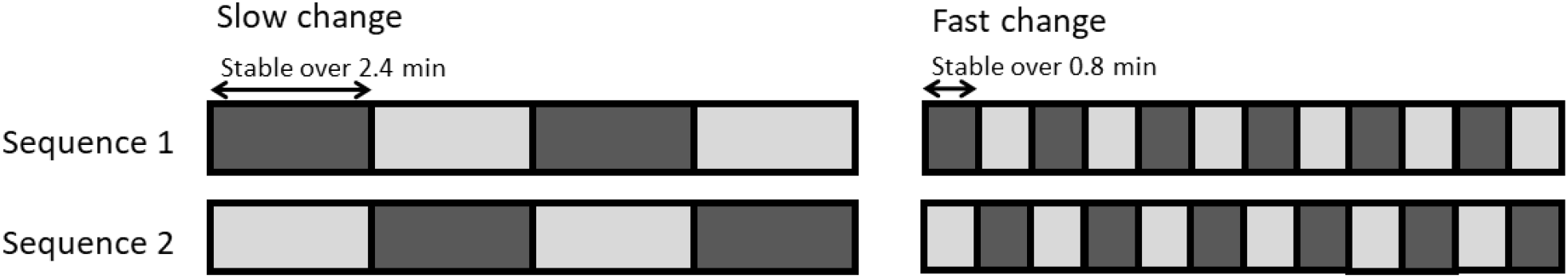
Structure of the multi-timescale MMN paradigm. Colored rectangles indicate the tone roles changing over blocks. In other words, the standard tone (p = 0.875) in dark grey blocks becomes the deviant tone (p = 0.125) in light gray blocks and vice versa. Roles in the initial block of the first sequence were counterbalanced across participants. Figure adapted from Todd and colleagues (2013).

In this paradigm, participants were presented via headphones with 1000-Hz pure tones, with durations of 60ms and 30ms at stimulus-onset asynchrony of 300ms. In each block, one of these tones was presented as the standard tone (probability of 0.875) and the other as the deviant tone (probability of 0.125). These roles alternated after every 480 tone presentations (2.4 minutes) in the slow change blocks, and after every 160 presentations (0.8 minutes) in the fast change blocks. A sequence consisted of four slow change blocks followed by 12 fast change blocks. Participants were presented with two sequences. In the second sequence, the roles in the initial block were opposite from that in the initial block of the first sequence. For example, if the 60ms tone was the deviant tone in the first block of the first sequence, the 30ms tone would take this role in the first block of the second sequence. Roles in the initial block of the first sequence were also counterbalanced across participants. Prior observations show that the effects are order-driven and not tone-driven (i.e., can be observed equally if the first deviant is a 30ms tone or a 60ms tone, Todd et al., 2013, 2020). There was a 1.5-minute break between the slow and fast blocks, and a 5-minute break between the two sequences, resulting in a total duration of approximately 47 minutes. All stimuli were presented binaurally with an intensity of 70dB through electroencephalogram (EEG)-compatible insert earphones (ER-3C, MedCaT, The Netherlands) using PsychoPy2 1.85.2. Participants were instructed to watch a silent, subtitled nature documentary and to ignore the sounds.

### Procedure

For participants with autism, the ADOS-2 and short form WAIS-III were completed during a first test session on a separate day. Four TD participants also completed the short form WAIS-III in a first session. The other TD participants were assessed with the short form WAIS-III at the end of the EEG session.

During the EEG session, participants first completed the multi-timescale MMN paradigm. After this, they took part in two more EEG experiments with an overall duration of approximately 30 minutes, of which the results will be presented elsewhere. Next, they filled in three questionnaires measuring autistic traits: the AQ (Baron-Cohen et al., 2001; Dutch version: Hoekstra et al., 2008), the SRS-A (Constantino, 2002; Dutch version: Noens et al., 2012), and the Adolescent/Adult Sensory Profile (SP; Brown & Dunn, 2002; Dutch version: Rietman, 2007).

### EEG Recordings and Analyses

EEG activity was recorded at a sample rate of 1024Hz using an ActiveTwo EEG amp (BioSemi) from 64 active Ag/AgCl scalp electrodes placed according to the international 10–20 system. Additional electrodes were applied at the mastoids, near the canthi and above and below the left eye. The data were referenced online to the common mode sense electrode. Data were recorded in an electrically shielded chamber. Electrode offsets were kept between −30 and 30µV at all electrodes.

EEG data analysis was performed using BrainVision Analyzer 2.1 and R 3.3.1 in RStudio 1.0.143, using the erpR package for plotting (Arcara & Petrova, 2014). Data were first rereferenced to the average of all scalp electrodes, downsampled to 500Hz, and filtered with a notch filter at 50Hz, a high-pass filter at 0.5Hz (12dB/oct), and a low-pass filter at 30Hz (12dB/oct). Trials were then epoched from 50ms before tone onset to 300ms after. Next, ocular artifacts were corrected with the Gratton and Coles method as implemented in BVA. Epochs exceeding an amplitude of ±75µV at the scalp or mastoid electrodes were rejected. When more than 10% of epochs were rejected due to amplitudes exceeding ±75µV at a specific electrode, this electrode was interpolated. A baseline correction was applied from −50 to 0ms relative to tone onset. Then, the data were rereferenced to the average of the mastoids (as recommended by Kujala et al., 2007). For two participants, we used only the left or right mastoid, due to excessive noise in the other mastoid. Next, 8 event-related potential (ERP) waveforms for deviant tones and 8 ERP waveforms for standard tones were created (i.e. for the 30ms and 60ms tones in both slow and fast blocks for each of the sequences). The first five trials of each block and the first standard after each deviant were excluded from averages (see also Todd et al., 2013). Finally, the ERPs were filtered with a 20Hz low-pass filter (12dB/oct), as recommended for MMN measurement (Kujala et al., 2007). As mentioned earlier, we excluded three participants in each group due to excessive alpha waves based on visual inspection of the individual average waveforms. For one participant, we interpolated an electrode in slow blocks of the first sequence only, based on visual inspection. There was no significant difference in the final number of trials between the TD and autism group, *t*(47.30)=-0.60, *p*=.55, see Supplemental Table 2 for number of trials per group (in each condition).

Difference waveforms (MMNs) for each condition were created by subtracting the waveform for the tone as a standard from the tone as a deviant. For example, the MMN for the 60ms tone in the slow blocks was created by subtracting the waveform for the 60ms tone when it was a standard in the slow blocks from the waveform for the 60ms when it was a deviant in the slow blocks (Jacobsen & Schröger, 2003). For statistical analyses, MMN amplitudes were then defined as the mean amplitude (± 20ms) surrounding the peak negativity between 100 and 220ms after stimulus onset in the difference waveforms, for each participant, electrode, and condition separately (similar to Goris et al., 2018). Based on earlier studies and the topography across groups, we included data of 3 frontocentral electrodes (F3, Fz and F4) (Baldeweg & Hirsch, 2015; Shaikh et al., 2012). A repeated-measures multivariate analysis of variance (MANOVA) was conducted on MMN amplitudes including group (2 levels: autism or TD) as a between-subject factor and initial role (2 levels: initial deviant and initial standard), speed (2 levels: slow and fast), sequence (2 levels: first and second) and electrode (3 levels: F3, Fz and F4) as within-subject factors.

For the block analyses reported below, MMN waveforms were created by grouping two blocks together, e.g. the MMN for the 60ms tone in the first two blocks of the slow blocks, was created by subtracting the waveform for this tone when it was a standard (e.g. block 1) from the waveform for when it was a deviant (e.g. block 2).

## Results

### Mismatch negativity

A complete overview of all effects in the repeated-measures MANOVA can be found in Supplemental Table 1. The mean MMN amplitudes for each group and each condition are presented in Figure 2. This figure shows clear group differences that seem to disappear for the second sequence, which was confirmed in the analysis.

**Figure 2.**
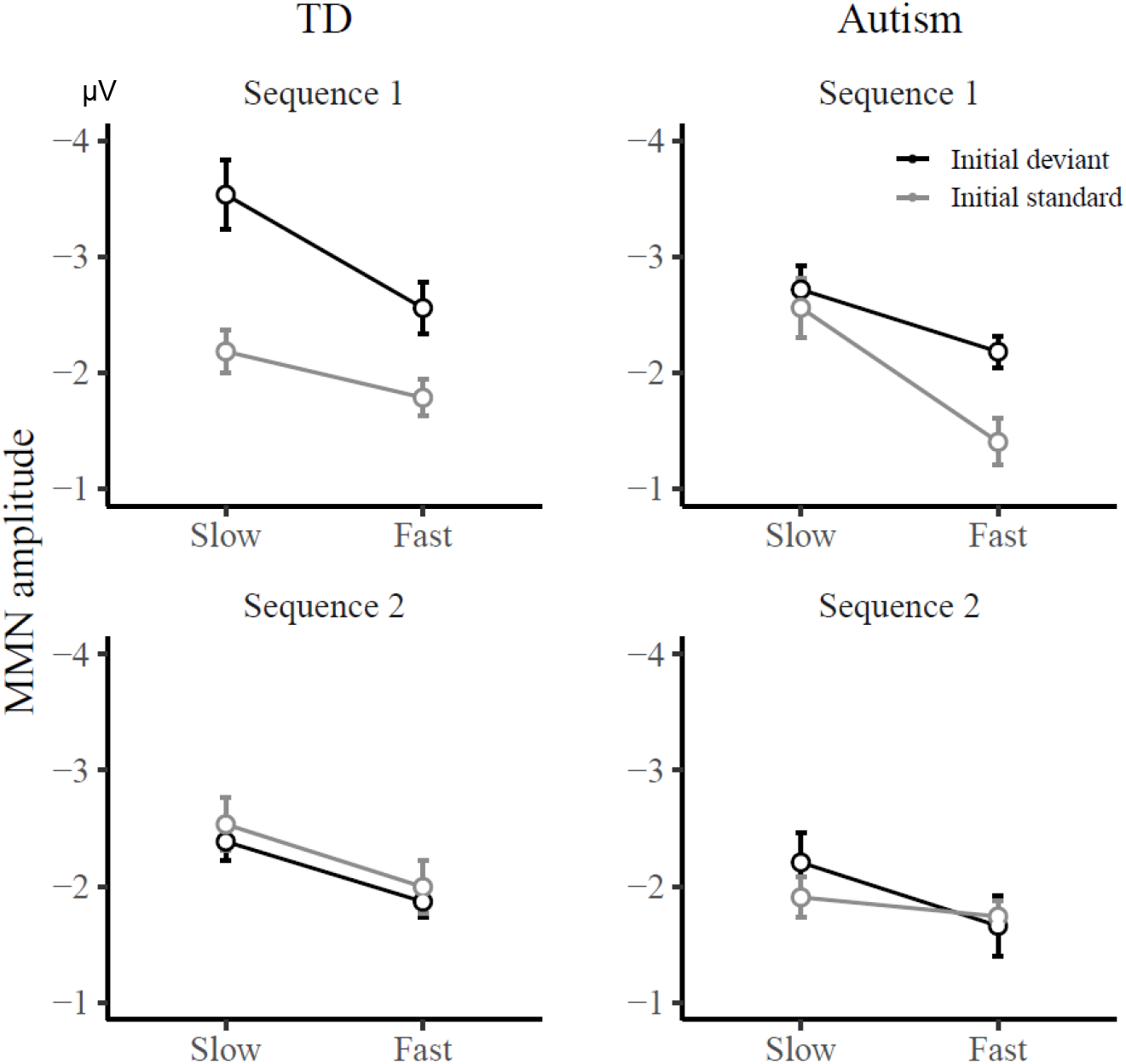
Group-averaged mismatch negativity (MMN) amplitudes to the initial deviant and initial standard, in the slow and fast speed condition for sequences 1 and 2, in the typically developed (TD) and autism group. Data were averaged over electrodes F3, Fz and F4. In the first sequence, there was a group by initial role interaction for the slow condition only. Specifically, in the TD group there was a difference in MMN amplitude between the initial deviant and initial standard, while this was not the case in the autism group.

The omnibus MANOVA showed a main effect of speed, *F*(1,52)=34.27, *p*<.001, *η*_*p*_^*2*^=0.40, indicating a larger MMN in slow versus fast changing blocks, suggesting that people, when given more time, build stronger models of what are deviant and what are standard sounds and hence show larger MMN amplitudes. Second, we found a main effect of initial role, *F*(1,52)=14.56, *p*<.001, *η*_*p*_^*2*^ = 0.22, indicating that participants show a larger MMN for the sound that was initially presented as the deviant sound, than for the sound that was initially the standard sound when it was later presented as a deviant. This suggests that participants have a stronger model when it is in line with the first experienced, primary context model, reflected in larger MMN amplitudes. Third, there was a main effect of sequence, *F*(1,52)=14.87, *p*<.001, *η*_*p*_^*2*^=0.22, showing that MMN responses decrease over time from the first to second sequence. We also found a significant 2-way interaction between group and electrode, *F*(2,51)=4.24, *p*=.02, *η*_*p*_^*2*^=0.14, suggesting a stronger effect of electrode in the TD compared to the autism group (see, Supplemental Figure 1). However, this main group difference in electrode did not interact with any other effect of interest. More importantly, previous studies with this design have shown an interaction between speed and initial role (Todd et al., 2011, 2013, 2014), which often further interacts with sequence (Todd et al., 2013), as a marker of how people update their model differently depending on whether it matches the first experienced, primary model or not – an observation that is usually most pronounced in the beginning of the experiment and can fade away over time (e.g., Todd et al., 2013, 2014). Across groups, neither this two-way interaction, nor three-way interaction, reached significance, both *F*(1,52)<0.35, both *p*>0.55.

Instead, the MANOVA showed a significant 4-way interaction between group, initial role, sequence and speed, *F*(1,52)=5.60, *p*=.02, *η*_*p*_^*2*^=0.10, in line with the idea that people with autism differ from TD subjects in how first impressions shape subsequent sensory processing. Follow-up tests for each sequence separately indicated a significant 3-way interaction between group, speed and initial role only in the first sequence, *F*(1,52)=9.22, *p*<.01, *η*_*p*_^*2*^=0.15, but not the second sequence, *F*(1,52)=0.75, *p*=.39, *η*_*p*_^*2*^=0.01, see Figure 2, suggesting that there were group differences in model updating, consistent with the idea of a differential primacy bias, during the first part of the experiment. In fact, the interaction between initial role and speed was not significant in the second sequence, *F*(1,52)=0.59, *p*=.44, *η*_*p*_^*2*^=0.01, see Figure 2, indicating that both groups no longer showed any sign of a primacy bias in the second half of the experiment. The Bayes Factor BF10 for the three-way interaction in the first sequence, however, was BF10=19.09, indicating strong evidence for the interaction (computed using Inclusion Bayes Factors across matched models as implemented in JASP version 0.10, JASP Team, 2019). Therefore, the remainder of the analyses focused on the first sequence only.

The three-way interaction between group, initial role, and speed, in the first sequence showed that people with autism differed from TD subjects, in how their MMN was modulated by the initial roles as well as the speed with which the tone probabilities changed (fast versus slow). In line with previous observations from Todd and colleagues, MMN amplitudes in the TD group were characterized by an initial role by speed interaction *F*(1,26)=3.35, *p*=.08, *η*_*p*_^*2*^=0.11. This pattern was reversed in the same interaction effect in the autism group, *F*(1,26)=6.87, *p*=.01, *η*_*p*_^*2*^=0.21.

To unpack the origins of these group differences, the slow and fast changing blocks were analyzed separately. Specifically, in the slow condition, there was a significant main effect of initial role for the TD group, *F*(1,26)=8.73, *p*<.01, *η*_*p*_^*2*^=0.25, meaning that MMN amplitudes were higher for initial deviants than for initial standards when they later become deviants, while the autism group showed no such difference between initial deviants and initial standards, *F*(1,26)=0.22, *p*=.64, *η*_*p*_^*2*^=0.01. These differences are clearly evident in Figure 2. This group difference in the effect of initial role in the slow condition was also supported by a group by initial role interaction in the slow condition, *F*(1,52)=4.45, *p*=.04, *η*_*p*_^*2*^=0.08. However, in the fast condition, both groups were comparable, *F*(1,52)=0.00, *p*=.99, *η*_*p*_^*2*^=0.00. This suggests that there was a primacy bias in the slow condition for the TD group, which was absent in individuals with autism. To investigate this result as a function of sequential learning, we further explored MMN amplitudes for each block in the first sequence separately.

### Analyses per block in the 1^st^ sequence

To enable a more serial inspection of the potential origin of group differences, separate difference waveforms were computed for each block encountered during sequence 1. As can be seen on Figure 4, the main group differences are evident in the slow condition blocks where the effect of initial role on MMN amplitudes could be observed across all four blocks in the TD group, suggesting it was not just an effect of the first block showing a larger MMN. When the initial deviant was again presented as a deviant in the third block, the MMN amplitude increased back to its original level. This suggests that participants were more inclined to rely on their initial model, and can thus be interpreted as a bias towards what participants encountered first. Importantly, this bias was completely absent in the autism group where the MMN amplitude was reasonably equivalent throughout. This was supported with a group by initial role interaction, *F*(1,52)=6.47, *p*=.01, *η*_*p*_^*2*^=0.11, indicating an effect of initial role only in the TD group, *F*(1,26)=9.42, *p*<.01, *η*_*p*_^*2*^=0.27, and not in the autism group, *F*(1,26)=0.01, *p*=.94, *η*_*p*_^*2*^=0.00. There were no main or interaction effects of block (grouped together by two blocks, resulting in a factor with 2 levels: the first two vs. the last two blocks). For the time course of MMN amplitudes within each block, see Supplemental Figure 3.

**Figure 3.**
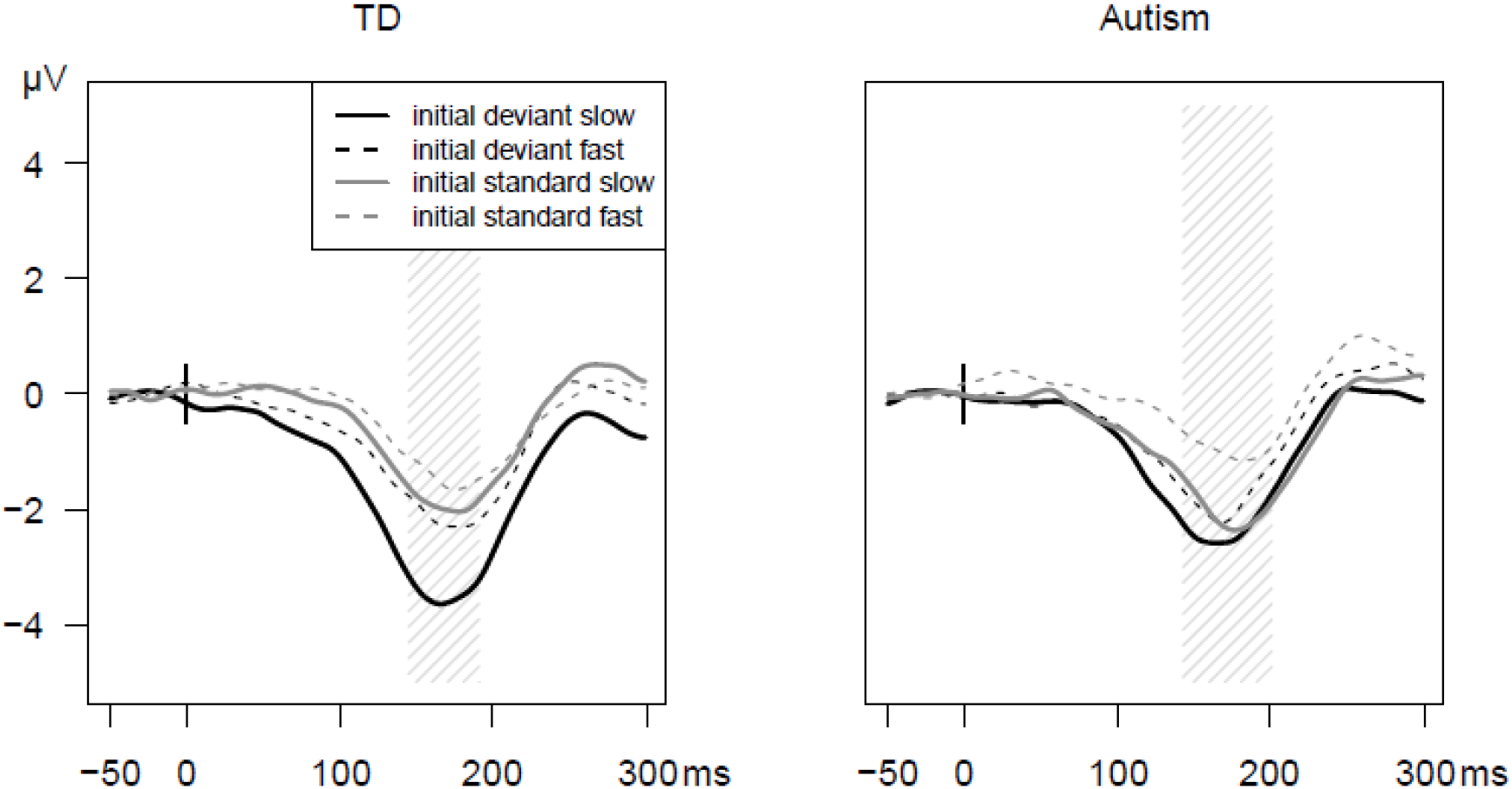
Group-averaged mismatch negativity (MMN) waveforms on electrode Fz in the first sequence only, for initial deviant and initial standard tones in the slow and fast speed condition, in the typically developed (TD) and autism group. The gray zone indicates 20 ms before until 20 ms after the mismatch negativity peaks. In the TD group, there was a difference between the initial deviant and the initial standard in the slow condition, while this was not the case in the autism group. As the MMN is the deviant minus standard difference waveform, in Supplemental Figure 2 we plotted waveforms in response to deviant tones (Supplemental Figure 2A) and in response to standard tones (Supplemental Figure 2B) separately.

**Figure 4.**
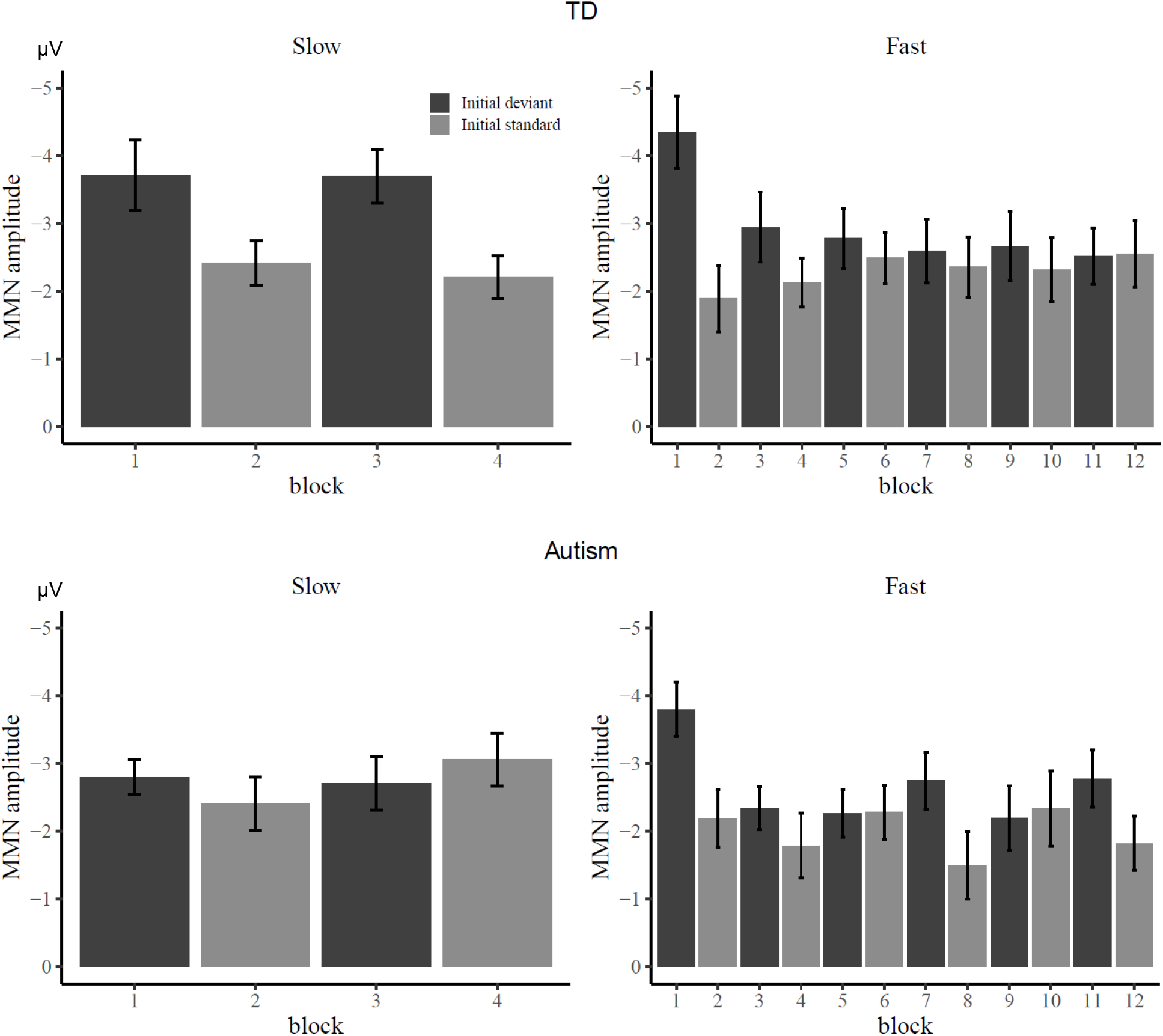
Group-averaged mismatch negativity (MMN) amplitudes, separately for each block of the slow and fast speed condition in sequence 1, in the typically developed (TD) and autism group. Data were averaged over electrodes F3, Fz and F4. Bar color indicates which tone was presented as the deviant tone in this block: the initial deviant tone (which was the deviant in the first block of the sequence), or the initial standard tone.

As groups did not differ in the fast condition, it is possible that either both groups do show a primacy bias here, or that there is an absence of this bias in both groups. Figure 4 suggests that the primacy bias disappears over time in the fast condition. Indeed, when investigating blocks separately (grouped together by two blocks, resulting in 6 levels), there was an initial role by block interaction, *F*(5,48)=3.51, *p*<.01, *η*_*p*_^*2*^=0.27, indicating a significant effect of initial role only in the first two blocks, *F*(1,52)=20.27, *p*<.001, *η*_*p*_^*2*^=0.28, but not in the other blocks. Therefore, rather than still reflecting a primacy bias, the remaining effect of initial role could be due to a heightened MMN after the short break between the slow and fast phase, as well as the unexpected fast reversal in contingencies in the second block of the fast phase. Consistent with our findings above, there were no interactions with group in the fast condition.

### Correlations between mismatch negativity and autistic traits

The above group analyses showed a clear group difference in primacy bias. In search for further convergent evidence, we also investigated whether this primacy bias effect was related to autistic traits as measured with questionnaires or ADOS-2 interview scores within these groups.

The speed by initial role interaction was quantified as the difference in MMN amplitude between the initial deviant and initial standard in the first slow condition, minus this difference in the first fast condition. This way, higher, positive numbers mean a stronger effect of initial roles in the slow condition compared to the fast condition. In both the TD and autism group, there were no significant correlations with the questionnaire scores, all |*r*|<.29, *p*>.14. However, when studying the ADOS-2 scores (which were only administered in the autism group), there was a marginally significant correlation between speed by initial role interaction and ADOS-2 total score, *r*=-.37, *p*=.06, see Figure 5A, suggesting that the interaction between speed and initial role might decrease with increasing autism symptom severity.

**Figure 5.**
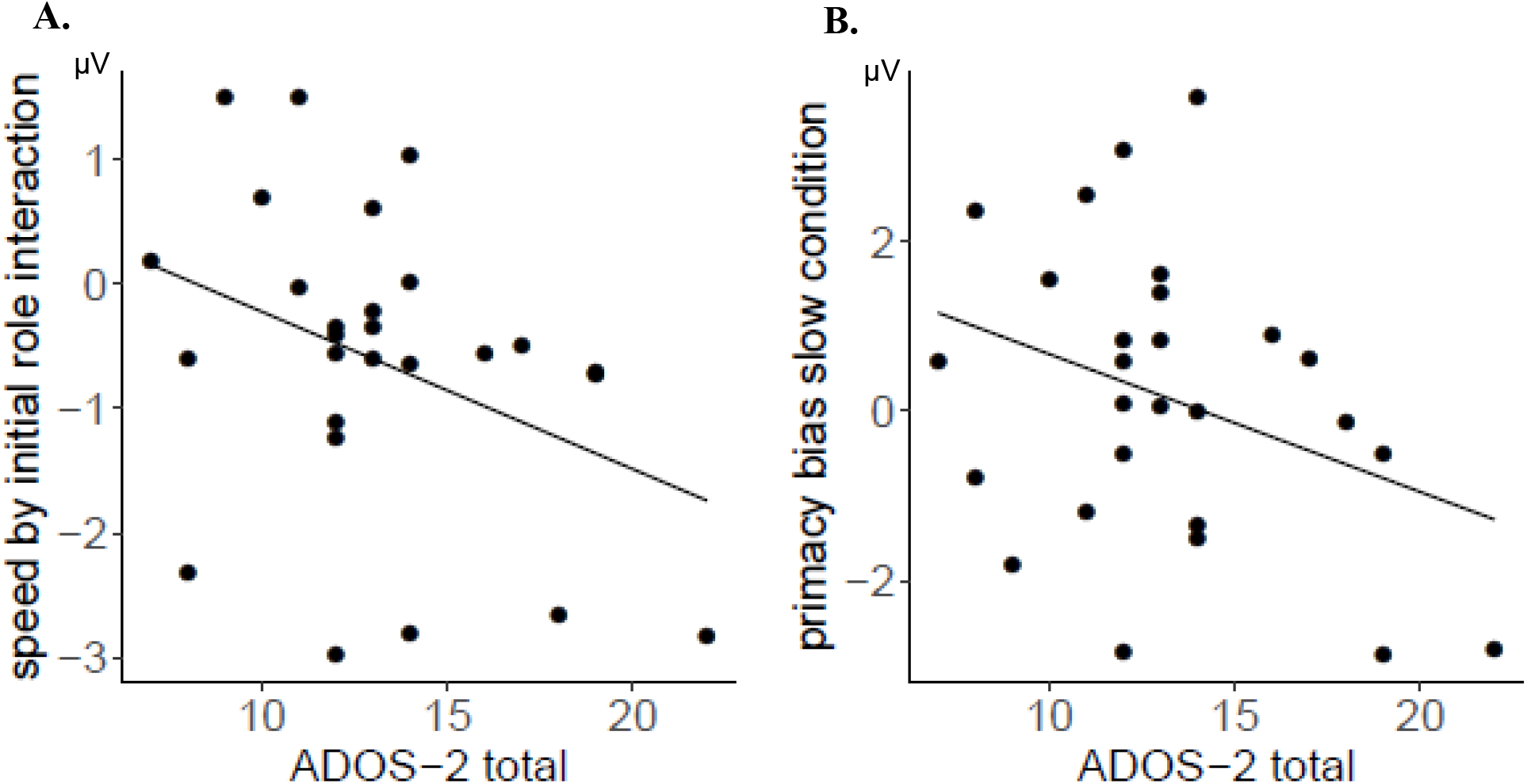
Scatter plots for the correlations between effects in mismatch negativity (MMN) amplitudes and the ADOS-2 total score in the ASD group. The trend lines reflect the Pearson correlation coefficients. (A) Correlation with the speed by initial role interaction in the first sequence. This interaction was quantified as the difference in MMN amplitude between the initial deviant and initial standard in the slow condition, minus this difference in the fast condition. (B) Correlation with the effect of initial role in the slow speed condition, i.e. the primacy bias. This was defined as the difference in MMN amplitude between initial deviants and initial standards in the slow condition of the first sequence.

Following up on this marginally significant correlation between the ADOS-2 score and the interaction effect, we also zoomed in on the correlations with the effect of initial role in the slow speed condition, i.e. the primacy bias, as this is where we previously found group differences. There was a marginally significant correlation, *r*=-.33, *p*=.09, see Figure 5B, hinting at a decreasing primacy bias with increasing autism symptom severity. There was no correlation with the effect of initial role in the fast condition, *r*=-.09, *p*=.67.

Summarized, these correlational results hint at a relation between autism symptomatology and reduced primacy bias. However, we have to be careful in interpreting this result, as this relation was only marginally significant and we did not correct for multiple comparisons.

## Discussion

Recent theories propose that autism characteristics can be understood from a predictive coding framework (Lawson et al., 2014; Van de Cruys et al., 2014; for a review see Palmer et al., 2017). They put forward that a core deficit in autism might be a high, context-insensitive weighting of prediction errors, which would lead to a faster incorporation of new information into modelled predictions in autism, compared with TD participants. On the other hand, slower model updating in autism has also been proposed (e.g. Lieder et al., 2019). In the current study, we set out to differentiate between these two hypotheses by investigating how models of the acoustical context are influenced by first impressions. We used the multi-timescale MMN paradigm by Todd and colleagues (2013), which has consistently shown that the initial model (i.e., which tone was the standard sound, and which the deviant, in a first series of trials) influences perception (reflected in MMN amplitudes) in later series of sounds, termed a *primacy bias*. Consistent with predictive coding theories, we expected that individuals with autism would be faster in updating models of sensory information and that this would be apparent as a smaller primacy bias (i.e. a reduced modulation of MMN amplitudes by the initial model). However, if autism is characterized by slower model updating, this should result in a larger primacy bias in MMN amplitudes.

Our results showed that, in the slow blocks presented at the beginning of the experiment, the TD group showed larger MMN amplitudes to initial deviant tones compared to initial standard tones when they were later presented as a deviant (consistent with earlier studies using a very similar design, Todd et al., 2011, 2014). This presumably indicates that TD participants confidently recognized the initial acoustic model when it re-emerged, resulting in more distinct responses for the deviant and standard tone and thus a larger MMN amplitude (which is defined as the difference between the responses to the deviant and standard tone). Crucially, this primacy bias was completely absent in the autism group, where we found no difference between the initial deviant and initial standard tone in the slow condition. This suggests that individuals with autism are not influenced by first impressions to the same extent as TD individuals, and can thus be interpreted as an indication of faster model updating (or abandoning of older information) in autism, consistent with predictive coding theories of autism.

In the subsequent fast blocks, there still seemed to be a difference between the initial deviant and initial standard for both groups. However, while we cannot fully exclude the idea that a primacy bias suddenly and surprisingly appears in the autism group there, we believe that this effect is likely due to increased attention after the break (leading to an increased MMN in the very first fast block), as well as the unexpected fast reversal in contingencies in the second block (presumably leading to a reduced MMN). Supporting this interpretation, the effect disappeared after these first two fast blocks for both groups, and is also absent in the second sequence. This is in line with earlier findings that show the primacy bias disappears over time (Todd et al., 2013).

Finally, interindividual difference analyses hinted at a correlation between our effects of interest and autism symptomatology in the autism group. While we have to interpret these effects with caution, this does support the idea that faster model updating is related to autistic characteristics. Specifically, we observed correlations with the ADOS-2 total scores only, and not with the questionnaires AQ or SRS-A. A possible explanation for this discrepancy might lie in the fact that the AQ and SRS-A are self-report questionnaires, while the ADOS-2 is an observational instrument.

Taken together, these results extend previous findings of reduced context-sensitivity of prediction errors in autism (Goris et al., 2018; Lawson et al., 2017; Palmer et al., 2015; Skewes et al., 2014) to not only support the hypothesis of a context-insensitive but also generally higher weighting of sensory prediction errors in autism (Lawson et al., 2014; Van de Cruys et al., 2014). In other words, our findings confirm the idea that new information is incorporated faster in sensory models in autism, and older information is abandoned faster, and demonstrate this at a very early level of sensory processing. Importantly, this weaker primacy bias found in sensory perception might not generalize to higher levels of information-processing. On the contrary, some studies have suggested a stronger primacy bias in autism during decision making (D’Cruz et al., 2013; Goris et al., 2019; Lieder et al., 2019; South et al., 2012). Possibly, people with autism overcompensate for faster model updating during early sensory processing, by being more conservative on higher levels of information processing. Future studies should systematically compare this seemingly divergent pattern of aberrant weighting of prediction errors during early sensory perception versus behavior at higher levels of information-processing (for example by investigating different types of primacy biases, e.g. Bronfman et al., 2016; Kotchoubey, 2014; Oberfeld et al., 2018). Another interesting research direction might be to formalize these observations and theories in a computational model of, for example, trial-by-trial MMN responses (e.g. Lieder et al., 2013) and P3b responses (Jepma et al., 2016; Kolossa et al., 2015).

In sum, we found that the MMN, an EEG component reflecting early sensory prediction errors, was less modulated by initial models in a group of adults with autism, compared to a control group, showing a weaker primacy bias in sensory perception in autism. This confirms the hypothesis of faster model updating, as put forward by predictive coding accounts of autism.

## Acknowledgements

J.G. was supported by a PhD fellowship and E.D. by a postdoctoral fellowship, both by the FWO – Research Foundation Flanders.

## Supplemental Material

**Supplemental Table 1.** Complete results for the repeated-measures analysis of variance exploring main and interactions effects on mean MMN amplitude to initial deviant and initial standard tones in slow and fast speed conditions for sequences 1 and 2, in the typically developed (TD) and autism group. Electrodes included were F3, Fz and F4. *** *p* < .001, * *p* < .05, . *p* < .10

| Effect | F-Statistic | P value |
| --- | --- | --- |
| Group | $F(1, 52) = 1.20$ | $p = .28$ |
| Sequence | $F(1, 52) = 14.87$ | $p < .001$ *** |
| Group x sequence | $F(1, 52) = 0.01$ | $p = .91$ |
| Speed | $F(1, 52) = 34.27$ | $p < .001$ *** |
| Group x speed | $F(1, 52) = 0.00$ | $p = .97$ |
| Initial role | $F(1, 52) = 14.56$ | $p < .001$ *** |
| Group x initial role | $F(1, 52) = 0.78$ | $p = .38$ |
| Electrode | $F(2, 51) = 15.78$ | $p < .001$ *** |
| Group x electrode | $F(2, 51) = 4.24$ | $p = .02$ * |
| Sequence x speed | $F(1, 52) = 3.56$ | $p = .06$ · |
| Group x sequence x speed | $F(1, 52) = 0.93$ | $p = .34$ |
| Sequence x initial role | $F(1, 52) = 3.58$ | $p = .06$ · |
| Group x sequence x initial role | $F(1, 52) = 1.05$ | $p = .31$ |
| Speed x initial role | $F(1, 52) = 0.35$ | $p = .56$ |
| Group x speed x initial role | $F(1, 52) = 2.19$ | $p = .14$ |
| Sequence x electrode | $F(2, 51) = 0.09$ | $p = .91$ |
| Group x sequence x electrode | $F(2, 51) = 0.88$ | $p = .42$ |
| Speed x electrode | $F(2, 51) = 0.48$ | $p = .62$ |
| Group x speed x electrode | $F(2, 51) = 1.31$ | $p = .28$ |
| Initial role x electrode | $F(2, 51) = 0.39$ | $p = .68$ |
| Group x initial role x electrode | $F(2, 51) = 0.90$ | $p = .41$ |
| Sequence x speed x initial role | $F(1, 52) = 0.35$ | $p = .56$ |
| Group x sequence x speed x initial role | $F(1, 52) = 5.60$ | $p = .02$ * |
| Sequence x speed x electrode | $F(2, 51) = 1.85$ | $p = .17$ |
| Group x sequence x speed x electrode | $F(2, 51) = 0.20$ | $p = .82$ |
| Sequence x initial role x electrode | $F(2, 51) = 0.20$ | $p = .82$ |
| Group x sequence x initial role x electrode | $F(2, 51) = 0.07$ | $p = .93$ |
| Speed x initial role x electrode | $F(2, 51) = 0.62$ | $p = .54$ |
| Group x speed x initial role x electrode | $F(2, 51) = 0.21$ | $p = .81$ |
| Sequence x speed x initial role x electrode | $F(2, 51) = 1.58$ | $p = .22$ |
| Group x sequence x speed x initial role x electrode | $F(2, 51) = 1.49$ | $p = .23$ |

**Supplemental Table 2.**
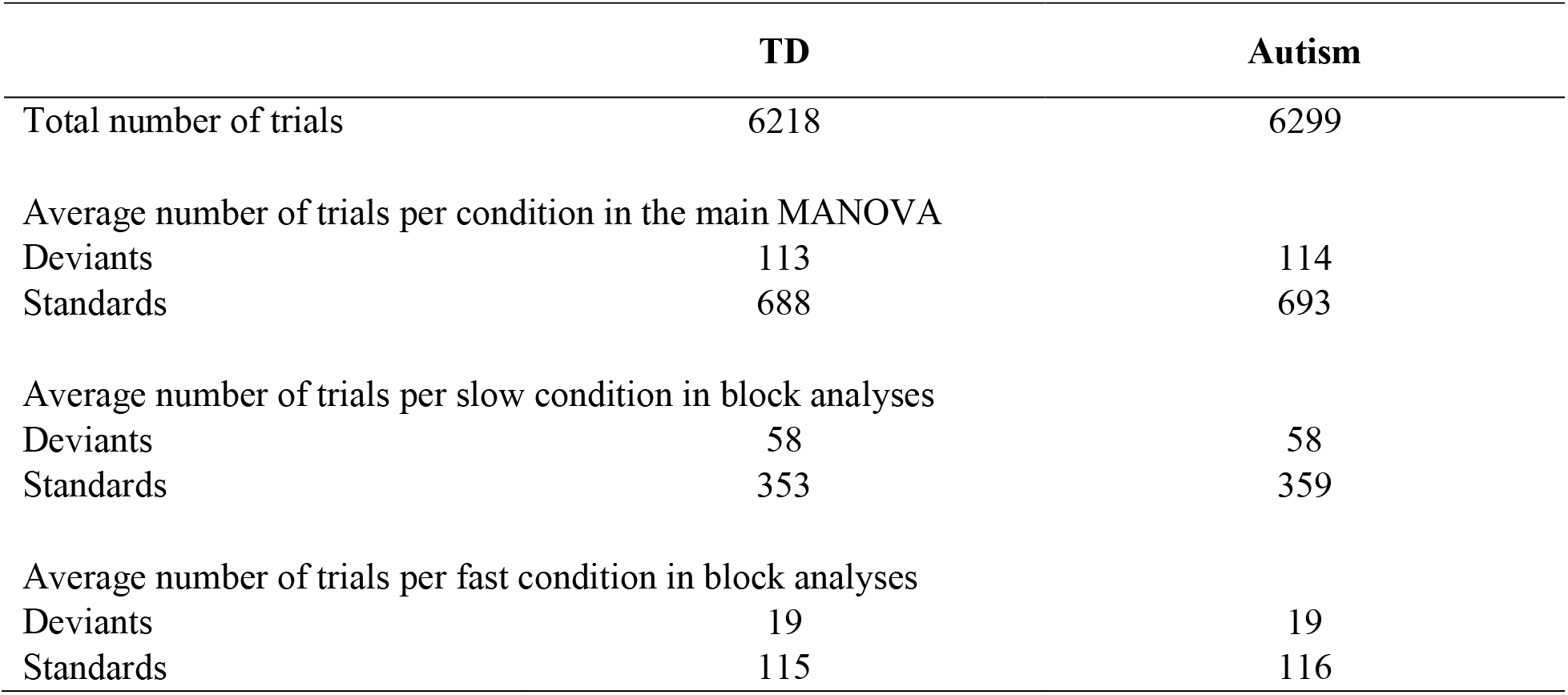
Average number of trials per condition, separately for the typically developed (TD) and autism group.

|  | **TD** | **Autism** |
| --- | --- | --- |
| Total number of trials | 6218 | 6299 |
| Average number of trials per condition in the main MANOVA |  |  |
| Deviants | 113 | 114 |
| Standards | 688 | 693 |
| Average number of trials per slow condition in block analyses |  |  |
| Deviants | 58 | 58 |
| Standards | 353 | 359 |
| Average number of trials per fast condition in block analyses |  |  |
| Deviants | 19 | 19 |
| Standards | 115 | 116 |

**Supplemental Figure 1.**
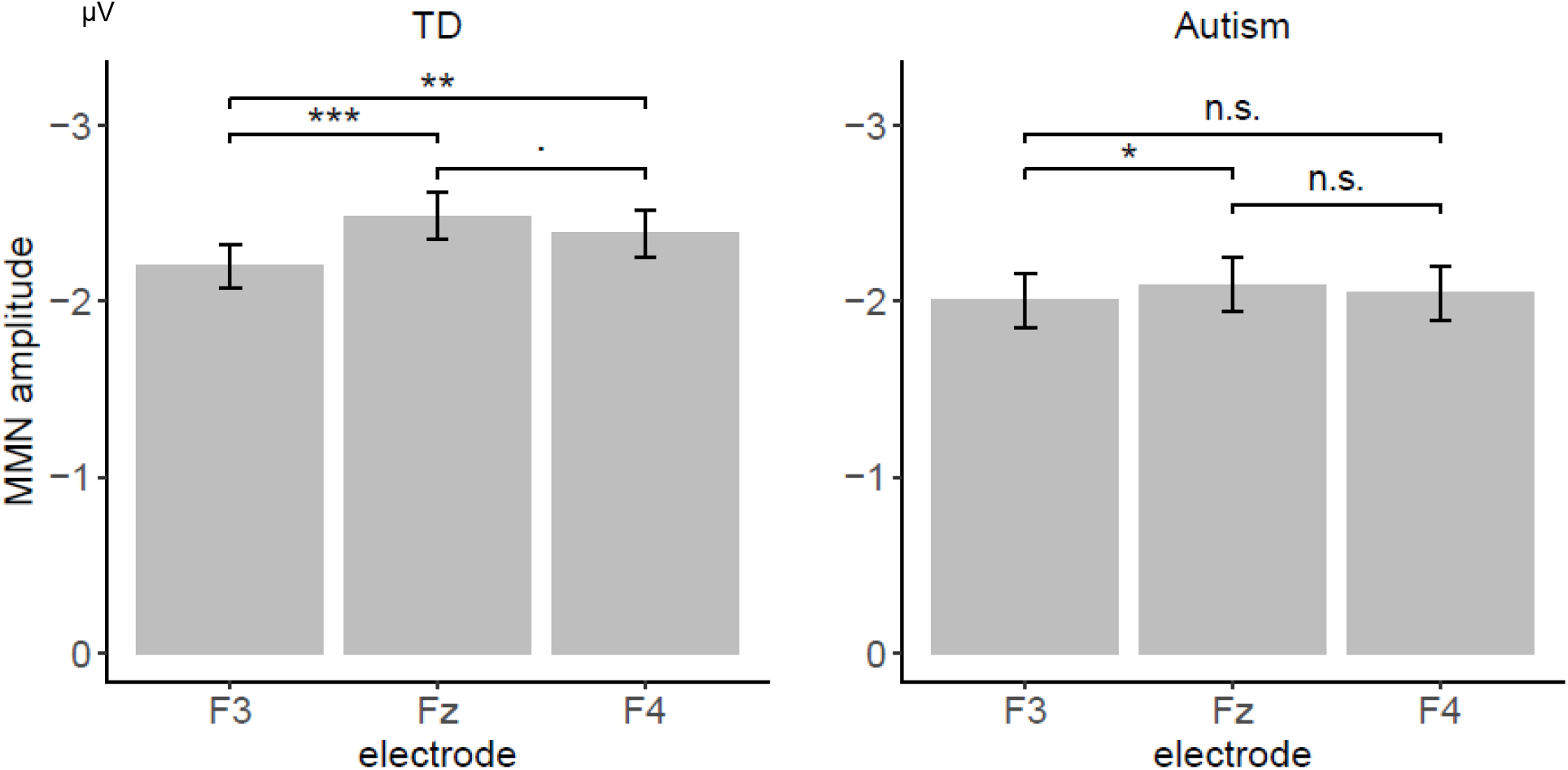
MMN amplitudes on electrodes F3, Fz and F4 in the typically developed (TD) and autism group. There was a significant interaction between group and electrode, F (2, 51) = 4.24, p = .02. This indicated a stronger effect of electrode in the TD group (F (2, 25) = 13.06, p < .001) than in the autism group (F (2, 25) = 2.85, p = .08). Follow-up t-tests in the TD group show a significant difference between the electrodes F3 and Fz, t (26) = −5.20, p < .001, as well as a marginally significant difference between Fz and F4, t (26) = 1.94, p = .06, and a significant difference between F3 and F4, t (26) = −2.97, p < .01. In the autism group, there was a significant difference between F3 and Fz, t (26) = −2.38, p = .02, but not between Fz and F4, t (26) = 0.79, p = .44, nor between F3 and F4, t (26) = −0.62, p = .54.

**Supplemental Figure 2.**
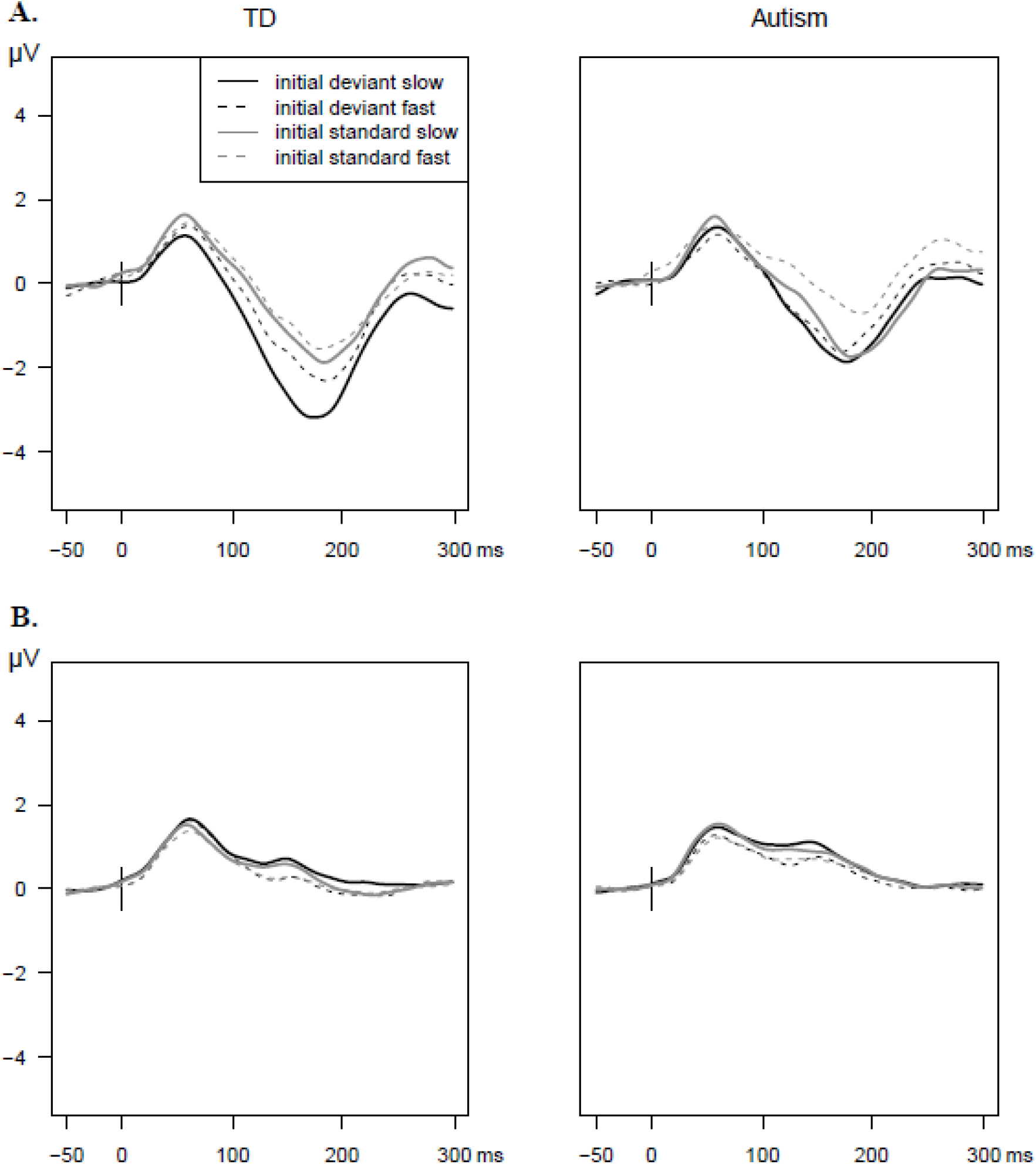
**(A)** Group-averaged ERP waveforms in response to *deviant* tones on electrode Fz in the first sequence only, for initial deviant and initial standard tones in the slow and fast speed condition, in the typically developed (TD) and autism group. **(B)** Group-averaged ERP waveforms in response to *standard* tones.

**Supplemental Figure 3.**
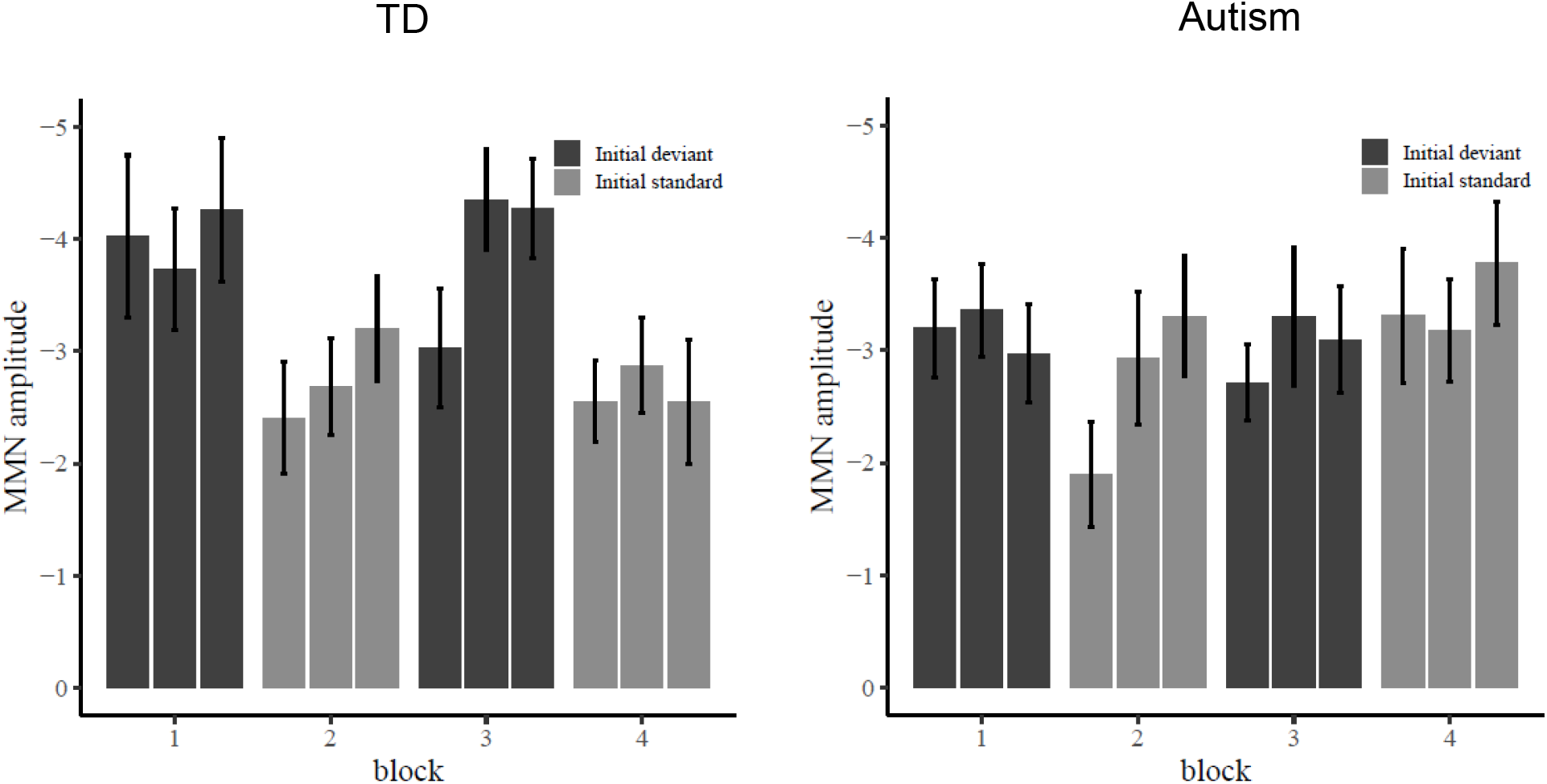
Group-averaged mismatch negativity (MMN) amplitudes for the slow blocks of the first sequence, each divided in three miniblocks, separately for the typically developed (TD) and autism group. MMN waveforms for each miniblock were created by grouping two blocks together, e.g. the MMN for the 60ms tone was created by subtracting the waveform for this tone when it was a standard (average over e.g. block 2) from the waveform for when it was a deviant (in one miniblock e.g. miniblock 1 of block 1). Average number of deviants per miniblock was *M*=19 for both the TD and autism group, average number of standards per block was *M*=353 for the TD group and *M*=359 for the autism group. Data were averaged over electrodes F3, Fz and F4. Bar color indicates which tone was presented as the deviant tone in this block: the initial deviant tone (which was the deviant in the first block of the sequence), or the initial standard tone.

